# Spatial and Temporal Localization of SPIRRIG and WAVE/SCAR Reveal Roles for These Proteins in Actin-Mediated Root Hair Development

**DOI:** 10.1101/2020.11.13.381848

**Authors:** Sabrina Chin, Taegun Kwon, Bibi Rafeiza Khan, J. Alan Sparks, Eileen L. Mallery, Daniel B. Szymanski, Elison B. Blancaflor

**Affiliations:** Noble Research Institute LLC, 2510 Sam Noble Parkway, Ardmore, Oklahoma 73401; Department of Botany and Plant Pathology, West Lafayette, Indiana 47907; Dept. of Biological Sciences, Purdue University, West Lafayette, Indiana 47907

## Abstract

Root hairs are single cell protrusions that enable roots to optimize nutrient and water acquisition. They attain their tubular shapes by confining growth to the cell apex, a process called tip growth. The actin cytoskeleton and endomembrane systems are essential for tip growth; however, little is known about how these cellular components coordinate their activities during this process. Here, we show that SPIRRIG (SPI), a BEACH domain-containing protein involved in membrane trafficking, and BRK1 and SCAR2, subunits of the WAVE/SCAR (W/SC) actin nucleating promoting complex, display polarized localizations to root hairs at distinct developmental stages. SPI accumulates at the root hair apex via post-Golgi vesicles and positively regulates tip growth by maintaining tip-focused vesicle secretion and filamentous-actin integrity. BRK1 and SCAR2 on the other hand, mark the root hair initiation domain to specify the position of root hair emergence. Consistent with the localization data, tip growth was reduced in *spi* and the position of root hair emergence was disrupted in *brk1 and scar1234*. BRK1 depletion coincided with SPI accumulation as root hairs transitioned from initiation to tip growth. Taken together, our work uncovers a role for SPI in facilitating actin-dependent root hair development through pathways that might intersect with W/SC.

## INTRODUCTION

Root hairs are single cell tubular projections that emerge from root epidermal cells. They increase the effective surface area of the root system by extending laterally into soil pores, thus enabling the increased access to nutrients and water (Carminati et al., 2017, Ruiz et al., 2020). Root hairs have been studied extensively by plant biologists for decades because they serve as excellent models to unravel mechanisms by which cell size and shape in plants is regulated (Grierson et al., 2014). To attain their cylindrical shapes, root hairs undergo tip growth, a process in which expansion of the cell is confined to its apical domain. Tip growth involves a balance between the directed delivery of post-Golgi vesicles carrying protein complexes and cell wall building blocks to the cell apex, and localized cell wall loosening and recycling of excess membranes. Besides root hairs, tip growth is exhibited by other cell types such as pollen tubes, fungal hyphae, and rhizoids of mosses, liverworts and algae (Bascom et al., 2018a).

Root epidermal cells called trichoblasts are the cell types that form root hairs. Work in *Arabidopsis thaliana* has shown that trichoblasts are specified to become root hair-forming cells early during root development through the patterned assembly of protein complexes of transcriptional activators and repressors in different cell files (Schiefelbein et al., 2014, Shibata et al., 2018). Upon establishing their identity, trichoblasts undergo two developmental stages that lead to root hair outgrowth. The first stage is the establishment of a root hair initiation domain (RHID) that eventually leads to a conspicuous root hair bulge at the basal (root-tip oriented) end of the trichoblast (Grierson et al., 2014, Nakamura and Grebe, 2018). Several proteins accumulate at the RHID, most prominently, the Rho of Plants (ROP)/RAC small GTPases and their Guanine Nucleotide Exchange Factor (GEF) activators (Denninger et al., 2019). The second stage is tip growth. In addition to small GTPases, the cytoskeleton, the endomembrane trafficking machinery, cytoplasmic calcium, phosphoinositide lipids, hormones (e.g. auxin and ethylene) and reactive oxygen species are other major players in signaling pathways that modulate root hair development (Bascom et al., 2018a, Nakamura and Grebe, 2018).

The filamentous-actin (F-actin) and microtubule cytoskeletons orchestrate root hair development. The actin cytoskeleton in particular has been studied widely during tip growth as it functions as tracks for the traffic of cellular cargo to the cell apex (Bascom et al., 2018a, Stephan, 2017, Szymanski and Staiger, 2018). Many insights into the role of the actin cytoskeleton in tip growth have been demonstrated through work with *Arabidopsis* mutants and pharmacological approaches involving the use of chemicals that disrupt F-actin. For example, the actin-disrupting compound, latrunculin B (LatB), inhibits tip growth, while also inducing the formation of root hairs and pollen tubes with irregular shapes (Bibikova et al., 1999, Gibbon et al., 1999). In this regard, *Arabidopsis* mutants with defects in the root hair-expressed vegetative *ACTIN2* (*ACT2*) gene are characterized by root hairs that mirror those treated with LatB (Ringli et al., 2002, Yoo et al., 2012). Furthermore, *ACT2* and *ACT7* mutants display an apical (shoot-ward) shift in the position of the RHID when compared to wild type (Kiefer et al., 2015).

To fulfill its cellular functions, the actin cytoskeleton is organized into higher order networks that correspond to the growth strategy of the cell. This is evident in tip-growing cells, whereby the base and shank of the cell consists mostly of thick, longitudinal F-actin bundles, while the apex contains actin fringes, rings, patches or a fine meshwork depending on the plant species or cell type (Stephan, 2017). To sustain tip growth, the integrity and organization of these tip-focused F-actin arrays must be maintained, a task that is facilitated by a diverse collection of actin-binding proteins and actin nucleators (Li et al., 2015, Paez-Garcia, 2018, Szymanski and Staiger, 2018).

The actin-related protein (ARP2/3) complex and its activator, suppressor of cAMP receptor (SCAR)-WASP family verprolin homologous (WAVE) complex (W/SC), is the best characterized actin nucleator in plants (Deeks and Hussey, 2005, Szymanski, 2005, Yanagisawa et al., 2013). Upon conversion from an inactive open conformation to an active closed conformation, the ARP2/3 complex promotes F-actin nucleation from the sides of existing filaments by forming a surface that mimics stable actin dimers (Blanchoin et al., 2000, Robinson et al., 2001, Rodal et al., 2005). Activation of ARP2/3 for efficient nucleation of F-actin requires the W/SC nucleation promoting factor (NPF) complex. In addition to W/SC, the NPF consists of the proteins SRA1, NAP1, ABI1 and HSPC300/BRICK1 (BRK1) (Basu et al., 2004a, Basu et al., 2005a, Djakovic et al., 2006, El-Assal et al., 2004, Zhang et al., 2005b, Zhang et al., 2008, Jörgens et al., 2010). Studies of *Arabidopsis* trichome development have been most instrumental in revealing insights into the function of W/SC-ARP2/3 in plants. Plant counterparts to the mammalian W/SC-ARP2/3 subunits were first uncovered through the cloning of disrupted genes in a set of *DISTORTED* (*DIS*) *Arabidopsis* trichome mutants (Hülskamp et al., 1994, Mathur et al., 2003, Le et al., 2003, El-Assal Sel et al., 2004, Basu et al., 2004b, Deeks et al., 2004, Zhang et al., 2005a). W/SC is the only known NPF for ARP2/3, and *BRK1* and *SCAR* mutants display null *arp2/3* trichome phenotypes (Le et al., 2006, Zhang et al., 2008).

In addition to trichome development, the W/SC-ARP2/3 actin filament-nucleating module has been implicated in specifying leaf pavement cell shape, light and auxin-dependent root growth, stomatal gating, gravitropism, responses to salinity stress, and plant immunity (Li et al., 2003, Basu et al., 2005b, Zhang et al., 2005a, Dyachok et al., 2011, Zhao et al., 2013, Li et al., 2014, Zou et al., 2016, Isner et al., 2017, Badet et al., 2019, Pratap Sahi et al., 2018). The W/SC andARP2/3 complexes has been shown to play an essential role in some tip growing cells such as the protonemal cells of the moss *Physcomitrella patens*. This was shown when RNA interference (RNAi) of transcripts encoding selected subunits of the moss W/SC andARP2/3 complexes resulted in substantial protonemal tip growth defects (Harries et al., 2005, Perroud and Quatrano, 2006, Finka et al., 2008). The importance of the W/SC andARP2/3 complexes was further demonstrated by Perroud and Quatrano (2006, 2008) who showed that two of its components, BRK1 and ARPC4, accumulated in tips of moss protonemal cells. By contrast, *Arabidopsis* plants with mutations in components of the W/SC andARP2/3 complexes displayed minimal or no defects in root hair and pollen tube growth (Li et al., 2003, Mathur et al., 2003), and none of the subunits have been conclusively shown to exhibit clear polar localization in these cell types. Therefore, the extent by which the W/SC andARP2/3 complexes functions in tip growing cells of higher plants remains to be determined.

*SPIRRIG* (*SPI*) is one of eight genes that has been considered to be a member of the *DIS* group, but compared to other *DIS* mutants, trichome phenotypes of *spi* are less severe and the mutant does not display early stage cell swelling that is diagnostic of the *DIS* group (Schwab et al., 2003). *SPI* was shown to encode a 3,571 amino acid long protein with N-terminally located armadillo (ARM) and concanavalin A (ConA)-like lectin domains and C-terminally-located pleckstrin homology (PH), beige and Chediak Higashi (BEACH) and WD40 repeat domains (Saedler et al., 2009). BEACH domain-containing proteins are highly conserved in eukaryotes and are known to function in membrane dynamics, vesicle transport, apoptosis and receptor signaling. This family of proteins are of clinical importance as they have been implicated in a variety of human disorders such as cancer, autoimmunity syndrome and autism (Cullinane et al., 2013).

In addition to mild trichome defects, *Arabidopsis SPI* mutants have short root hairs characterized by fragmented vacuoles suggesting that SPI, like other eukaryotic BEACH domain-containing proteins, functions in membrane trafficking (Saedler et al., 2009). Additionally, an observation by Steffens et al. (2017) that showed SPI physically interacting with proteins involved in endosomal sorting reinforces its role in membrane remodeling. SPI was also demonstrated to localize in mRNA processing bodies (P-bodies) in transiently transfected *Arabidopsis* epidermal cells, thereby suggesting a novel role for SPI in post-transcriptional regulation (Steffens et al., 2015). Because *SPI* had a substantial but incomplete overlap with distorted mutant phenotypes, it was suggested that SPI might be involved in actin-mediated cell developmental processes (Saedler et al., 2009), and perhaps function in coordination with W/SC and ARP2/3 complexes. However, SPI is not a known W/SC or ARP2/3 subunit; therefore, its relationship to ARP2/3 function is unclear. Moreover, the uncertainty with regard to SPI function is confounded by the fact that its subcellular localization in root hairs, which exhibit the most profound phenotype when *SPI* is mutated, remains unknown. In this paper, we addressed these questions by generating a fully functional SPI fluorescent protein fusion, and documented its spatial and temporal localization during root hair development. In addition, we used live cell microscopy to simultaneously image the dynamics of SPI and W/SC, and correlate SPI localization with root hair tip-focused F-actin. Taken together, our results suggest potential functional relationships among SPI, W/SC and actin during root hair development.

## RESULTS

### Isolation of a New *SPIRRIG* Mutant Allele

We previously described a forward genetic screen that led to the isolation of three non-allelic recessive *Arabidopsis* mutants that were hypersensitive to the growth inhibitory effects of LatB. The *hypersensitive to LatB1* (*hlb1*) and *hlb3* mutants have been described previously (Sparks et al., 2016, Sun et al., 2019). *hlb1* was disrupted in a gene encoding a *trans*-Golgi Network-localized tetratricopeptide repeat protein involved in actin-mediated membrane recycling (Sparks et al., 2016), whereas the genetic lesion in *hlb3* was found to encode the class II actin nucleator formin (Sun et al., 2019). Here, we report on *hlb2*, the third of these recessive mutants. Like *hlb1* and *hlb3*, *hlb2* primary root growth was more severely inhibited by LatB when compared to wild type. In the absence of LatB or at low (i.e. 25 nM) LatB concentrations, the primary root length of *hlb2* was similar to wild type. Differences in root length between wild type and *hlb2* became apparent when seedlings were grown on a concentration of 50 nM LatB and higher (Supplemental Figure 1A to C).

Through map-based cloning, we found that the mutation in *hlb2* was confined to a region between the *AT1G02740* and *AT1G03410* loci. Nucleotide sequencing revealed that *hlb2* had a 10 base pair deletion (Chr1 position: 720,152-720,161) in exon 14 of the *AT1G03060* gene. This 10-base pair deletion led to an open-reading frame shift at the Asp^1526^ codon resulting in a truncated protein (Supplemental Figure 2A). *AT1G03060* encodes the BEACH domain-containing protein, SPI (Saedler et al., 2009) (Supplemental Figure 2A). Because the first *spi* mutant alleles were reported to have short root hairs (Saedler et al., 2009), we examined root hairs of *hlb2*. We found that *hlb2* root hair growth rate was about 70% slower than wild type with some root hairs forming only small bulges (Supplemental Figure 2B and C). Moreover, *hlb2* had mild trichome defects that were reminiscent of the phenotypes of previously isolated *spi* mutant alleles (Supplemental Figure 2D; Saedler et al., 2009). To further verify if *HLB2* is *SPI*, we obtained a mutant from the publicly available SALK collection (SALK_065311), which had a T-DNA insertion in the 10^th^ exon of the *SPI* gene (Supplemental Figure 2A; Alonso et al. (2003)). The SALK_065311 line we obtained is the same as *spi-3* mutant allele reported previously in Steffens et al. (2015). In addition to having similar root hair and trichome defects as *hlb2*, primary roots of *spi-3* were hypersensitive to LatB (Supplemental Figure 2E), and a cross between *hlb2* and SALK_065311 failed to complement each other in the F1 hybrid. Taken together, these results indicate that *hlb2* is a new *spi* mutant allele. Based on earlier nomenclature (Steffens et al., 2015), we renamed *hlb2* to *spi-5* (Supplemental Figure 2A). The *spi-5* mutant allele was used for all subsequent experiments.

### SPIRRIG Localizes to the Tips of Rapidly Elongating Root Hairs

SPI fluorescent protein fusions have been previously demonstrated to associate with P-bodies and endosomes (Steffens et al., 2015, Steffens et al., 2017). However, these constructs have not been shown to complement the defective root hair and trichome phenotypes of *spi*. Because of the large size of SPI, we were unable to generate native promoter-driven fluorescent protein fusions to the full length *SPI* complementary DNA or genomic DNA. To overcome this problem, SPI was tagged with the 3x-Yellow Fluorescent Protein (<u>Y</u>F<u>P</u>) for <u>e</u>nergy transfer (YPet; Nguyen and Daugherty (2005)) using a method called recombineering. This method involves tagging the gene of interest in the context of transformation-competent bacterial artificial chromosomes to ensure that all regulatory sequences are included in the fluorescent protein fusion (Brumos et al., 2020, Zhou et al., 2011).

Once *spi* was transformed with a recombineered *SPI-YPet* construct, primary root hypersensitivity to LatB, short root hairs and defective trichome phenotypes were rescued, indicating that the construct was functional (Figure 1A; Supplemental Figure 3). Transgenic complementation of *spi* with *SPI-YPet* provided additional evidence that *HLB2* is *SPI*. Confocal microscopy of more than 150 root hairs from at least 25 seedlings revealed that SPI-YPet signal was strongest in the tips of rapidly elongating root hairs (Figure 1B and C; Supplemental Movie 1 and 2). Low magnification time lapse movies showed that 100% of rapidly elongating root hairs had robust SPI-YPet signal at the tips (Supplemental Movie 1). SPI-YPet fluorescence was not detected or weak at the RHID and during early root hair bulge formation, but intensified as root hairs transitioned to tip growth (Figure 1C; Supplemental Movie 3). SPI-YPet signal dissipated as root hairs matured and tip growth rate declined (Figure 1B and C; Supplemental Movie 4). A linear regression analysis showed that the intensity of SPI-YPet fluorescence in root hair tips is significantly positively correlated to rapid root hair growth (Figure 1D). These results indicate that the SPI protein has functions related to root hair elongation and consistent with the short root hair phenotypes of *spi*.

**Figure 1.**
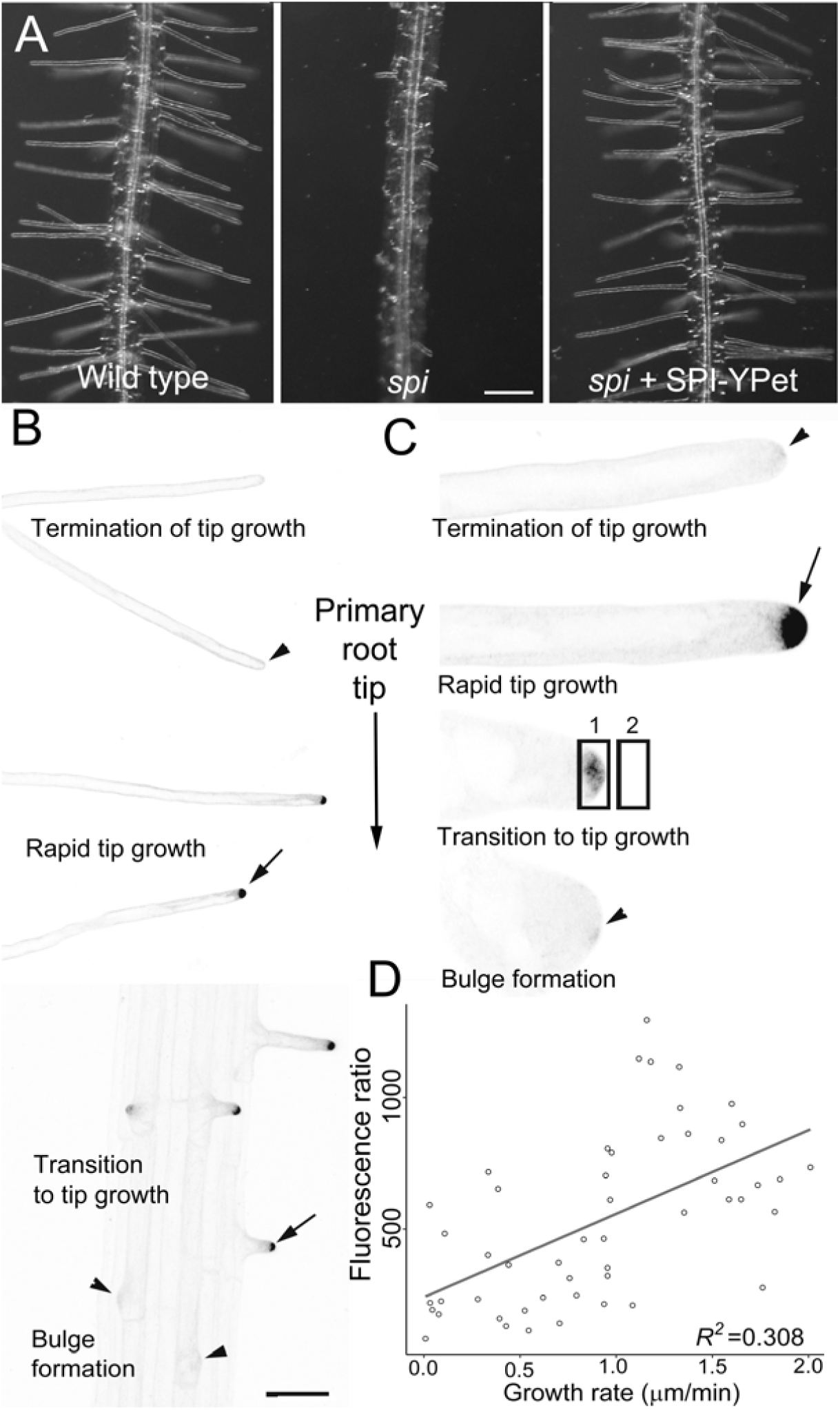
A Functional SPI-YPet Fusion Localizes to the Tips of Growing Root Hairs. **(A)** SPI-YPet rescues the short root hair phenotype of *spi*. Bar = 100 μm. **(B)** Low magnification image shows that SPI-YPet signal is most prominent at the tips of rapidly expanding root hairs (arrows). A black and white look up table (LUT) image is provided to increase clarity of SPI-YPet localization. SPI-YPet signal (black color) is weak during initiation/early bulge formation and mature root hairs that have terminated tip growth (arrowheads). Bar = 50 μm **(C)** High magnification images of single root hairs during bulge formation until tip growth termination. A black and white look up table (LUT) image is provided to increase clarity of SPI-YPet localization. SPI-YPet (black color) is enriched at the tip of root hairs that are rapidly growing or transitioning to tip growth (arrow). Faint SPI-YPet signal is found in bulging root hairs or those that have stopped elongating (arrowheads). Images are representative of about 150 root hairs from at least 25 seedlings. Bar = 10 μm. **(D)** Scatter plot showing correlation analysis of root hair tip SPI-YPet fluorescence and root hair tip growth. The mean fluorescence in the oval in region 1 divided by the oval in region 2 as shown in panel **C** represents the fluorescence ratio in the Y axis. Line shows linear regression fit with *R^2^* value = 0.308 and *p*= 1.72 x 10^-5^. (n=5-7 root hairs per time point).

### SPIRRIG is Transported to the Root Hair Tip via Post-Golgi Vesicles

The prominent SPI-YPet signal at the tips of elongating root hairs is reminiscent of the localization patterns of post-Golgi markers such as RAB small GTPases, which are known to function in tip-directed secretion (Preuss et al., 2004). We therefore hypothesized that SPI is trafficked to the tips of root hairs via post-Golgi vesicles. To test this hypothesis, seedlings expressing *SPI-YPet* were treated with brefeldin A (BFA). BFA is a fungal toxin that is routinely used to investigate endomembrane dynamics because it prevents vesicle formation for exocytosis by inhibiting ADP ribosylation factor guanine nucleotide exchange factors (ARF-GEFs), while still enabling endocytosis and some retrograde pathways to continue (Baluška, 2002, Doyle et al., 2015). Consequently, BFA treatment causes the formation of endosomal agglomerations called BFA bodies. Treatment of seedlings expressing *SPI-YPet* with 50 μM BFA induced the formation of SPI-YPet agglomerates in root hairs (Figures 2A to C). The average number of BFA bodies per root hair was 3.69 ± 4.05 (S.D). Quantification of the BFA effect was also conducted by obtaining the fluorescence ratio of the SPI-YPet agglomerates to the fluorescence of the root hair cytoplasm that did not contain any agglomerates (Figure 2D, inset). The higher fluorescence ratio of BFA-treated root hairs compared to untreated controls reinforces our qualitative observations of the sensitivity of SPI-YPet to BFA (Figure 2D). Appearance of fluorescent puncta at the shank of root hairs also suggest that SPI-YPet is trafficked to the tip via post-Golgi vesicles (Supplemental Movies 2-4). These results indicate that SPI-YPet is associated with endomembranes and is trafficked to the root hair tips via BFA-sensitive post-Golgi vesicles.

**Figure 2.**
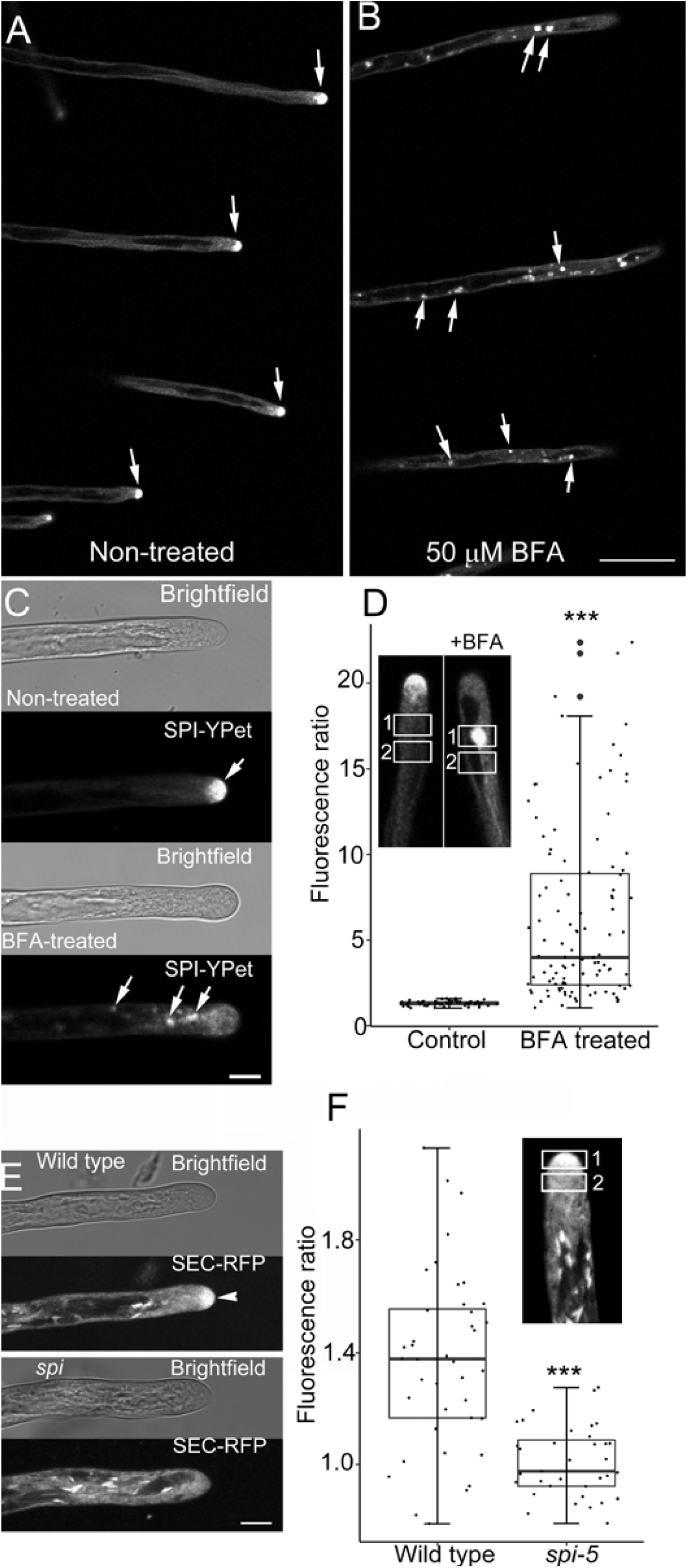
SPI-YPet is Localized to Brefeldin A (BFA) Sensitive Post-Golgi Compartments. **(A)** to **(B)** Low magnification image showing several root hairs expressing SPI-YPet. Note that untreated root hairs maintain tip-focused SPI-YPet while those treated with BFA show an abundance of fluorescence agglomerates (arrows). Bar = 50 μm. **(C)** Bright field and corresponding fluorescence image of representative untreated and BFA-treated elongating root hair showing the accumulation of SPI-YPet at the apical dome in solvent control-treated seedlings (arrow). Within 10 min after treatment with 50 μM BFA, SPI-YPet at the root hair tip dissipated and formed fluorescent agglomerates along the subapical regions (arrows). Bar = 10 μm. **(D)** Box plot of BFA induced agglomerates of SPI-YPet in untreated control root hairs and after treatment with 50 μM BFA. Ratio values were obtained by dividing mean fluorescence in rectangular region in 1 over region 2 (inset). Box limits indicate 25^th^ and75^th^ percentiles, horizontal line is the median and whiskers display minimum and maximum values. Each dot represents individual measurements from 8-14 root hairs per group from 8-24 plants. Asterisk (***) indicates statistical significance (*p*<0.001) as determined by Student’s T-test. BFA treated plants had an average of 3.7 BFA-induced agglomerates per root hair, with standard deviation of 4.05, whereas control root hairs showed no BFA-induced agglomerates per root hair. **(E)** The bulk secretory marker, SEC-RFP, accumulates at the tips of growing wild-type root hairs (arrowhead), but is absent in *spi* root hairs. Bar = 10 μm. **(F)** Box plot of SEC-RFP root hair tip accumulation expressed as fluorescence ratio. Ratio values were obtained by dividing mean fluorescence in oval region in 1 over region 2 (inset). Box limits indicate 25^th^ and 75^th^ percentiles, horizontal line is the median and whiskers display minimum and maximum values. Asterisk (***) indicates statistical significance (*p*<0.001) as determined by Student’s T-test. Each dot represents individual measurement from 8-10 root hairs per group from 9-12 plants

Slow root hair growth of *spi* and the formation of SPI-YPet agglomerates after BFA treatment, suggest that *spi* may be defective in tip-directed protein secretion and partly explain the short root hair phenotypes of *spi*. To address this question, we expressed the secreted (SEC)-red fluorescent protein (RFP) that contains a cleavable sporamin signal peptide that accumulates in the apoplast (Faso et al., 2009) and observed changes in tip-directed exocytosis using the membrane selective dye FM 1-43 dye in both *spi* and wild type genotypes (Bolte et al., 2004, Jelínková et al., 2010, Malínská et al., 2014). Similar to previous reports, rapidly elongating wild-type root hairs had a tip-focused gradient of SEC-RFP (Sparks et al., 2016) (Figure 2E). By contrast, slow-growing root hairs of *spi* lacked these tip-focused SEC-RFP gradients (Figure 2E). Tip-focused SEC-RFP in elongating wild-type and *spi* root hairs was quantified by obtaining the ratio of tip fluorescence to the subapical cytoplasm (Figure 2F, inset). The lower fluorescence ratio of *spi* root hairs compared to wild type confirmed our qualitative observations (Figure 2F). These results were corroborated with FM 1-43 dye uptake results, in which *spi* root hairs showed significantly reduced tip focused FM 1-43 gradient (Supplemental Figure 4A to C). Altogether, the loss of tip-focused secretion indicated defects in tip-directed bulk flow exocytosis in *spi* mutants.

### SPIRRIG Maintains Root Hair Tip-Focused F-actin

In a study of other *spi* mutant alleles, the similarities in phenotypes between *spi,w/sc andarp2/3* mutants suggests that SPI could function in actin-dependent cellular processes (Saedler et al., 2009). However, because trichome defects of *spi* were mild compared to other *w/sc* and *arp2/3* mutants, no obvious actin phenotypes were observed in *spi* trichomes (Schwab et al., 2003). To clarify the relationship between SPI and actin, we focused on investigating actin organization in root hairs since they displayed the most obvious growth defects in *SPI*-altered plants.

To study actin organization, we expressed the live F-actin reporter, *UBQ10: mGFP-Lifeact*, in *spi* (Vidali et al., 2010). This particular F-actin reporter was selected because it prominently labels the tip-focused F-actin meshwork typically observed in root hairs that are rapidly growing (Sparks et al., 2016). In wild type, fine F-actin networks were observed in the root hair bulge that had a weaker signal compared to the thicker actin bundles in other regions of the trichoblast (Figure 3A). As the root hair bulge expanded and the root hair transitioned to rapid tip growth, the tip-focused F-actin meshwork, which consisted of short filaments and dynamic puncta, became more conspicuous (Figures 3B-E; Supplemental Movie 5; Sparks et al. (2016)). When wild-type root hairs stopped elongating, the tip-focused F-actin meshwork was replaced with thick F-actin cables (Figure 3F).

**Figure 3.**
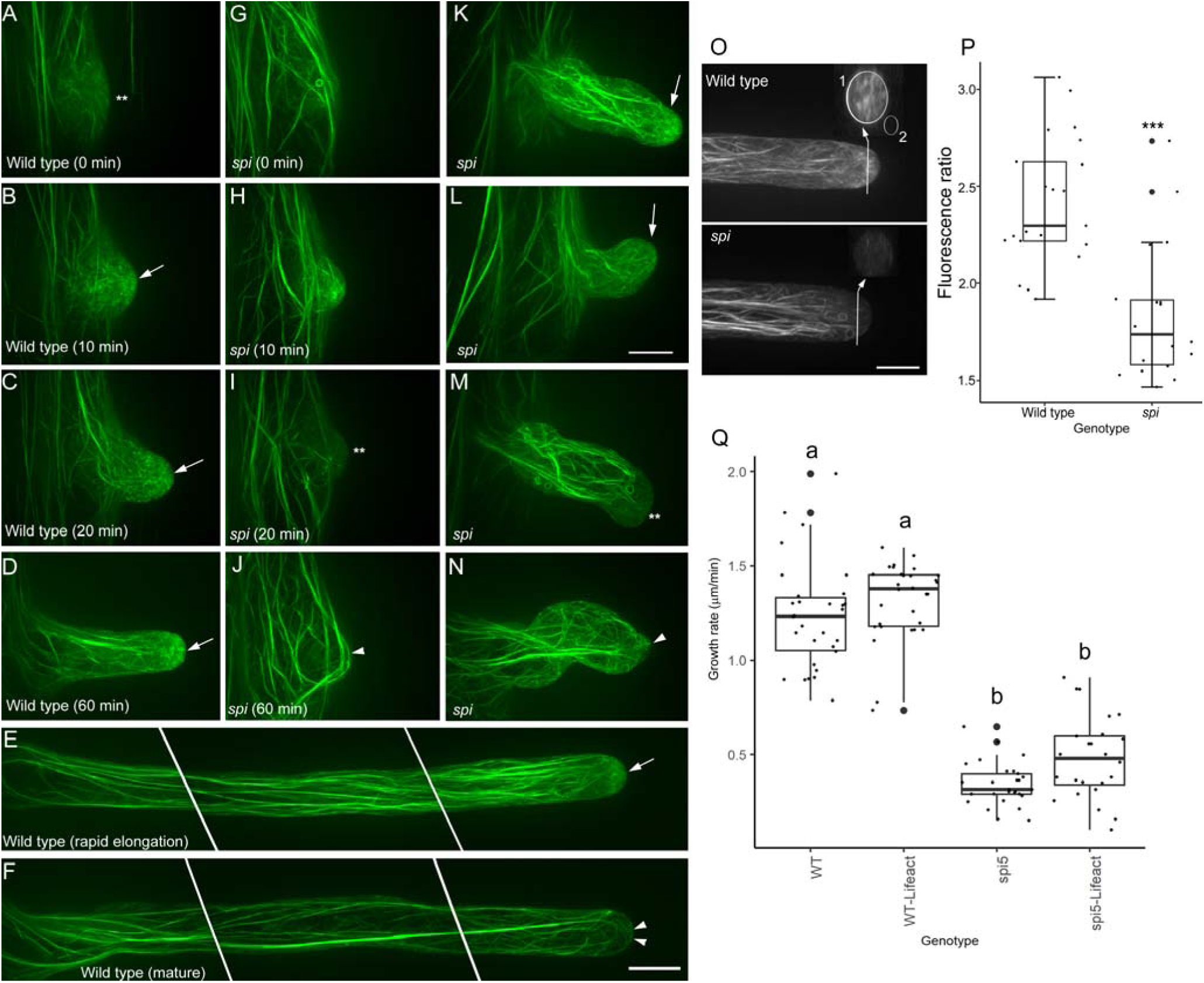
The Tip-focused F-actin Meshwork is Disrupted in Root Hairs of *spi*. **(A)** to **(D)** Time course of F-actin organization in a wild-type root hair from bulge formation to rapid tip growth. Weakly fluorescing F-actin networks (**) in the root hair bulge **(A)** reorganize into prominent tip-focused meshworks as the root hair transitions to rapid tip growth (arrows in **B** to **D**). **(E)** Tip-focused F-actin meshworks (arrow) remain prominent in a long, rapidly elongating wild-type root hair. **(F)** Tip-focused F-actin meshworks in wild-type root hairs are replaced with F-actin bundles (arrowheads) that protrude to the tip when growth stops. **(G)** to **(J)** Time course of F-actin organization in a *spi* root hair bulge that is unable to transition to tip growth. Distinct F-actin meshworks are unable to form in root hair bulges that terminate tip growth (**, **I**). Thick F-actin bundles eventually form in these short, non-growing root hair bulges (arrowhead, **J**). **(K)** to **(N)** F-actin organization in slow-growing *spi* root hairs. Some *spi* root hairs show the tip-focused F-actin meshworks typically observed in wild type (arrow, **K, L**). However, tip-focused F-actin meshworks in slow-growing *spi* root hairs either dissipate (**, **M**) or prematurely form thick F-actin bundles that protrude to the tip (arrow, **N**). Images from **(A)** to **(N)** are based on maximum projection images of 20-25 optical sections taken at 0.5 μm intervals. **(O)** Representative maximum projection images and corresponding computer-generated cross-sections of wild-type and *spi* root hair tips. Only growing root hairs with a clear cytoplasmic cap were selected for analysis. The fluorescence ratio of the root hair tip (oval in 1) to background (oval in 2) was used to quantify tip-focused F-actin meshworks. **(P)** Box plot showing tip-focused F-actin fluorescence ratio. Box limits indicate 25^th^ and 75^th^ percentiles, horizontal line is the median and whiskers display minimum and maximum values. Asterisk (***) indicates statistical significance (*p*<0.001) as determined by Student’s T-test. Each dot represents individual measurement from 18-21 root hairs per group from at least 5 independent seedlings. **(Q)** Comparison of root hair growth rates between wild type and *spi* lines with their corresponding live F-actin reporter lines,*UBQ10: mGFP-Lifeact*,. Box limits indicate 25^th^ and 75^th^ percentiles, horizontal line is the median and whiskers display minimum and maximum values. Letters indicates statistical significance (*p*< 0.05) as determined by one-way ANOVA. Each dot represents individual measurement from 4-5 root hairs per group from 1-2 plants.

F-actin organization in the tips of *spi* root hairs was different from that of wild type. As noted, some root hairs of *spi* were only able to form small bulges due to premature termination of tip-growth (Supplemental Figure 2). In *spi* root hairs, the distinct F-actin meshwork observed in wild-type root hairs was unable to form. Instead, F-actin in these *spi* root hair bulges contained thick F-actin cables that resembled those of wild-type root hairs that had terminated growth (Figure 3G and H). However, the thick F-actin cables in non-growing *spi* root hair bulges were unstable as they dissipated (Figure 3I) and reformed again at a later time (Figure 3J). In the small population of *spi* root hairs that were able to undergo tip growth, a few exhibited the tip-focused F-actin meshwork that resembled those observed in elongating wild-type root hairs (Figure 3K). However, most of these slow growing *spi* root hairs lacked the tip-focused F-actin meshwork (Figure 3L and M; Supplemental Movie 5) or had thick F-actin bundles protruding to the tip, a feature that was reminiscent of non-growing, mature wild-type root hairs (Figure 3N). The disruption of the tip-focused F-actin meshwork in *spi* was quantified by measuring F-actin fluorescence from computer reconstructed transverse sections of the root hair tip and by taking the ratio of the average fluorescence over background signal. A higher ratio indicates a higher signal of tip-focused F-actin (Figure 3O). Our analysis showed that the fluorescence ratio in *spi* root hairs was significantly reduced compared to wild-type root hairs, supporting visual observations that the tip-focused F-actin meshwork in *spi* root hairs is disrupted (Figure 3P).

To ensure that the mGFP-Lifectact probe did not interfere with normal root hair elongation or F-actin organization, we compared the growth rates of wild type and *spi* with and without the reporter. The root hair growth rates of wild-type root hairs without the mGFP-Lifeact probe was not significantly different from wild-type root hairs expressing the reporter. Similarly, the growth rate of *spi* expressing mGFP-Lifectact was not significantly different from *spi* without the reporter (Figure 3Q).

We next generated plants expressing both *SPI-YPet* and *mRuby-Lifeact* so we could correlate SPI and F-actin in growing root hairs. In elongating root hairs of dual labeled seedlings, SPI-YPet and the mRuby-labeled F-actin meshwork overlapped at the tip of actively elongating root hairs (Figure 4A; Supplemental Movie 6). In one time lapse sequence, when the root hair stopped growing at the 80 – 120 min time points, both SPI-YPet and mRuby-labeled F-actin meshwork dissipated from the root tip (Figure 4A). Quantification of both SPI-YPet and mRuby-Lifeact from at least three root hair time lapse sequences revealed that the appearance of both markers at the tip are highly correlated to each other (Figure 4B and C). This supports that root hair tip-localized SPI is strongly associated with the tip-focused F-actin meshwork, and as such is involved in sustaining normal root hair elongation in coordination with actin.

**Figure 4.**
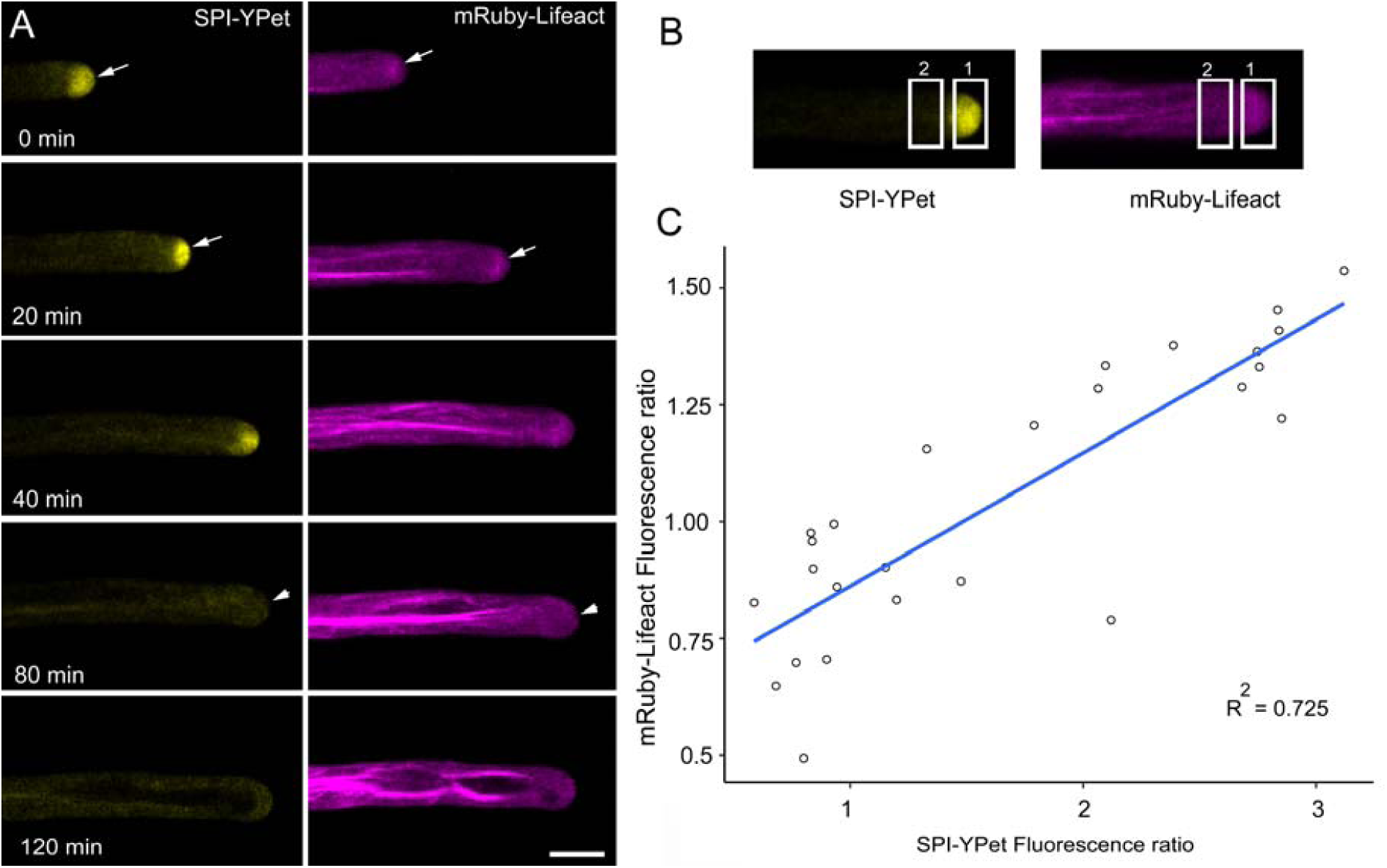
SPI-YPet and mRuby Lifeact Co-localizes at the Root tip in Elongating Root Hairs. **(A)** Time course of a root hair simultaneously expressing SPI-YPet and mRuby-Lifeact. Note that SPI-positive post-Golgi compartments and F-actin meshworks colocalized at the root hair apex (arrows) and dissipated at around the same time (arrowheads) at 80 min. Images are single median optical sections. Bars = 10 μm. **(B)** Method for obtaining SPI-YPet and mRuby-Lifeact ratios at the root hair tip for data shown in **C.** A rectangular region of interest at the tip and sub apex was used to measure fluorescence. **(C)** Scatter plot showing correlation analysis of root hair tip mRuby-Lifeact fluorescence and SPI-YPet fluorescence within the same root hair. The mean fluorescence in the rectangle in region 1 divided by the rectangle in region 2 as shown in panel **C** represents the fluorescence ratio for each reporter. For each ratio value, root hair growth rate was obtained by measuring the displacement of the root hair tip after a 10 min interval. Line shows linear regression fit with *R^2^* value = 0.725 and *p*= 2.127 x 10^-8^. (n = 26 time points from 3 root hair sequences)

### BRK1 and SCAR2 are Molecular Determinants of the Root Hair Initiation Domain

The identification of *SPI* as one of the genes in the *DIS* group that also included genes encoding subunits of the W/SC and ARP2/3 complexes (Saedler et al., 2009) raises the possibility that SPI might function in root hair developmental pathways mediated by W/SC-ARP2/3. Furthermore, the observation that SPI-YPET accumulation at the root hair tip (Figure 1) mirrors the enrichment of BRK1-YFP and ARPC4-GFP at apex of *Physcomitrella* protonemal cells (Perroud and Quatrano, 2006, Perroud and Quatrano, 2008) raises the possibility that W/SC and ARP2/3 complexes may be a root hair tip-localized complex. In an attempt to link SPI with the W/SC and ARP2/3 pathways, we imaged roots of *brk1* complemented with *BRK1promoter:BRK1-YFP* (from here on referred to as BRK1-YFP) (Figures 5A-C) (Dyachok et al., 2008). Unlike SPI-YPet, we did not observe a BRK1-YFP fluorescence gradient in rapidly elongating root hairs. However, closer examination of trichoblasts revealed prominent BRK1-YFP signal at the plasma membrane of the RHID that mirrored other known early root hair initiation site markers such as ROP (Figure 5A). Unlike ROP, which had a persistent plasma membrane localization throughout root hair development (Jones et al., 2002, Molendijk et al., 2001), BRK1-YFP signal was transient and dissipated as root hairs transitioned to rapid tip growth (Figure 5B; Supplemental Movie 7). This observation was in contrast to the intensification of SPI-YPet fluorescence as root hairs proceeded with rapid tip growth (see Figure 1). In agreement with our visual observations, a linear regression analysis showed that BRK1-YFP fluorescence is inversely proportional to root hair growth rate (Figure 5C).

**Figure 5.**
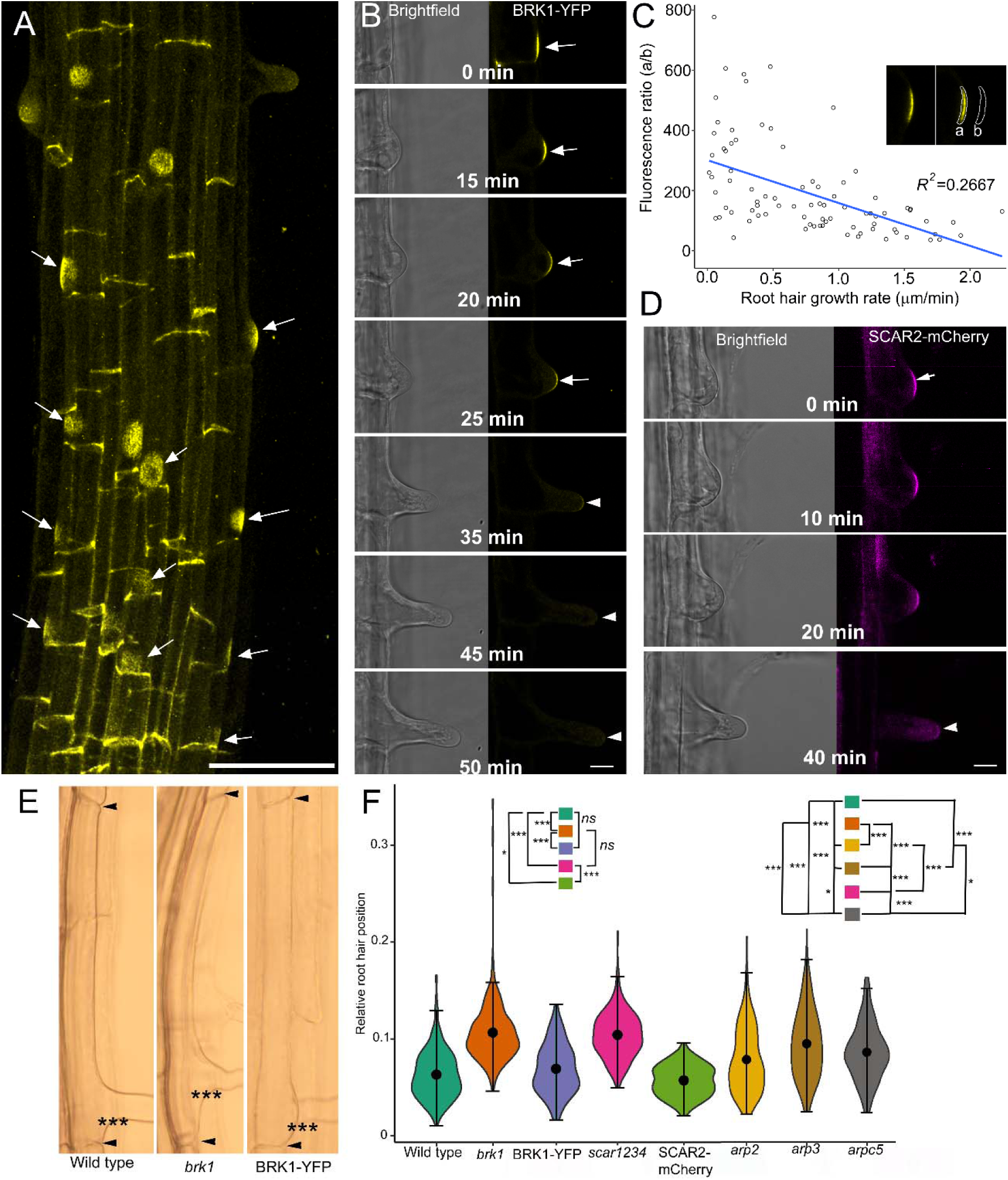
BRK1 and SCAR2 Mark the Root Hair Initiation Domain and Contribute to Planar Polarity. **(A)** Maximum projection confocal micrograph of the elongation and maturation zone of an *Arabidopsis* primary root expressing a functional BRK1-YFP fusion. The image was generated by merging 50 Z sections taken at 0.5 μm intervals. BRK1-YFP accumulates in the basal end walls and root hair initiation domains (arrows). Bar = 50 μm. **(B)** Time course of BRK1-YFP depletion in a developing root hair. BRK1-YFP signal (arrows) is strongest prior to the formation of a root hair bulge (0 min) and gradually dissipates (arrowheads) as the root hair undergoes rapid tip growth (35-50 min). Bar = 20 μm. **(C)** A scatter plot showing an inverse relationship between BRK1-YFP signal and root hair growth rate. Ratio of fluorescence of BRK1-YFP (a) to background (b) (inset) was plotted against root hair growth rate. Line shows linear regression fit with *R^2^* value = 0.2667 and *p*= 1.131 x 10^-7^. (n= 5-7 root hairs per time point). **(D)** Time course of SCAR2-mCherry in a developing root hair. Like BRK1-YFP, SCAR2-mCherry signal is strongest at the RHID and early stages of root hair bulge formation (0 min, arrow). SCAR2-mCherry signal dissipates when the root hair undergoes rapid tip growth (40 min, arrowhead). Bar = 20 μm. **(E)** Brightfield microscopy images of representative trichoblasts from 6-d-old seedlings showing apical shift in root hair position of *brk1* compared to wild type, and complementation of *brk1* planar polarity phenotypes by BRK1-YFP. Arrowheads mark the end walls of the trichoblast and asterisks (***) mark the basal wall of the emerged root hair. **(F)** Violin plots of root hair planar polarity in wild type, *brk1, scar1234, BRK1-YFP* in *brk1*, *SCAR2-mCherry* in *scar1234, arp2, arp3* and *arpc5* genotypes. Relative root hair position was obtained by taking the ratio of the distance from the basal trichoblast wall (bottom arrowheads in **E**) to the basal root hair wall (***) over the total length of the trichoblasts (i.e. length between the two arrowsheads). The plot illustrates kernel probability density in which the width represents distribution of data points. The black dot is the median and whiskers display minimum and maximum values. Statistical significance was determined using non-parametric, two sample Kolmogorov-Smirnov (KS) pairwise test. Wild type versus *brk1* (\*\*\**p* < 2.2 x 10^-16^); *brk1* versus BRK1-YFP in *brk1* (\*\*\**p*< 2.2 x 10^-16^); Wild type versus BRK1-YFP (*p*=0.0808 not significant, *ns*); Wild type versus *scar1234* (\*\*\**p* < 2.2 x 10^-16^); *brk1* versus *scar1234* (*p*= 0.699, *ns*); *scar1234* versus SCAR2-mCherry in *scar1234* (\*\*\**p* < 2.2 x 10^-16^); Wild type versus *SCAR2-mCherry* in *scar1234* (\**p*=0.012); Wild type versus *arp2* (\*\*\**p=* 2.739 x 10^-5^); Wild type versus *arp3* (\*\*\**p*= 8.882 x 10^-16^); Wild type versus *arpc5* (\*\*\**p*= 9.18 x 10^-12^); *brk1* versus *arp2* (\*\*\**p*< 2.2 x 10^-16^) *; brk1* versus *arp3* (\*\*\**p*= 3.41 x 10^-8^)*; brk1* versus *arpc5* (\*\*\**p*= 1.304 x 10^-11^), *arp2* versus *arp3* (\*\*\**p=* 0.00036)*; arp2* versus *arpc5* (\**p*= 0.001624)*; arp3* versus *arpc5* (* *p*= 0.0465)*; brk1* versus *arp2*(\*\*\**p* < 2.2 x 10^-16^); *brk1* versus *arp3* (\*\*\**p* = 3.41 x 10^-8^); *brk1* versus *arpc5* (\*\*\**p* = 1.304 x 10^-11^). n = 90-117 root hairs.

Given that BRK1 stabilizes the entire family of SCAR proteins and is required for functional W/SC assembly (Le et al., 2006), we investigated if the SCAR protein localized to the RHID, similarly to BRK1. To address this question, we imaged a recombineered SCAR2-mCherry fusion expressed in the *scar1 scar2 scar3 scar4* (*scar1234*) quadruple mutant. We found that like BRK1-YFP, SCAR2-mCherry marked the RHID and dissipated when rapid root hair tip growth commenced (Figure 5D).

The accumulation of BRK1-YFP and SCAR2-mCherry at the RHID led us to hypothesize that *brk1* and *scar1234* might have defects in root hair initiation. One parameter that has been studied extensively as an indicator of root hair initiation defects is planar polarity, which is a measure of root hair position along the length of the trichoblast (Nakamura and Grebe, 2018). We found that root hair position of *brk1* and *scar1234* shifted apically (i.e. toward the shoot) when compared to wild type (Figure 5F). The planar polarity defects of *brk1* was rescued by expressing *BRK1-YFP* in *brk1* while partial complementation of *scar1234* was achieved by expressing *SCAR2-mCherry* in *scar1234* (Figure 5F).

We also investigated if the ARP2/3 pathway was involved in RHID since W/SC complex can interact with the ARP2/3 pathway. APR2/3 subunit mutants, *arp2, arp3* and *arpc5* demonstrated significantly apically shifted root hairs that were similar to *brk1* and *scar1234* (Figure 5F). Nonetheless, we did not observe polarized accumulation of ARPC5-GFP at RHID nor root hair tips (Supplemental Figure 5). This indicates that ARP2/3 pathway may interact with W/SC complex at RHID, but is not essential for root hair positioning.

Taken together, our results revealed that BRK1 and SCAR2 are new molecular determinants of the RHID and are required for specifying the position of root hair emergence and can do so in both ARP2/3 dependent and independent manner.

### SPI is Required for the Depletion of BRK1 as Root Hairs Transition to Tip Growth

Live cell microscopy of BRK1-YFP and SPI-YPet revealed contrasting spatial and temporal dynamics, with the former intensifying and the latter dissipating as root hairs elongated (Figure 1 and 5). To observe BRK1 and SPI simultaneously within the same root hair, we generated plants expressing both *SPI-YPet* and *BRK1-mRuby3*. Similar to BRK1-YFP, BRK1-mRuby3 labeled the RHID and dissipated as the root hair bulge expanded (Figure 6A). Within the same root hair cell, SPI-YPet fluorescence at the tip intensified as BRK1-mRuby3 dissipated (Figure 6A; Supplemental Movie 8). To better understand the relationship between SPI and BRK1, we expressed *BRK1-YFP* in *spi* and *SPI-YPet* in *brk1*. We found that unlike BRK1-YFP in the complemented *brk1* background (Figure 5B), BRK1-YFP signal in *spi* persisted throughout the entire imaging time course (Figure 6B). In several cases, BRK1-YFP remained visible in non-growing root hair bulges of *spi* for more than 60 min. The persistence of BRK1-YFP signal was also observed in short root hair outgrowths of *spi* (Figures 6C; Supplemental Movie 9). To support our visual observations with quantitative data, we selected root hairs of *spi* and *brk1* expressing BRK1-YFP that were of about equal lengths. From these root hairs, the ratio of BRK1-YFP tip fluorescence to subapical fluorescence was obtained (Figure 6D). These analyses showed that BRK1-YFP signal persisted in *spi* as demonstrated by the higher fluorescence ratio (Figure 6E). By contrast, BRK1-YFP in *brk1* root hairs disappeared from RHID when root hairs experienced rapid tip growth and thus did not exhibit fluorescence gradient at the root tips (Figures 6D and E). In addition, SPI-YPet in *brk1* exhibited similar dynamics as SPI-YPet in the complemented *spi* (Figure 6F). In *brk1*, SPI-YPet signal was weak during early root hair bulge formation and intensified as root hair tip growth accelerated (Figure 6F). Taken together, these results strongly indicate that SPI contributed to the stability of BRK1 during the transition from root hair initiation to rapid tip growth. On the other hand, BRK1 did not appear to have direct effects on SPI during root hair initiation and rapid tip growth.

**Figure 6.**
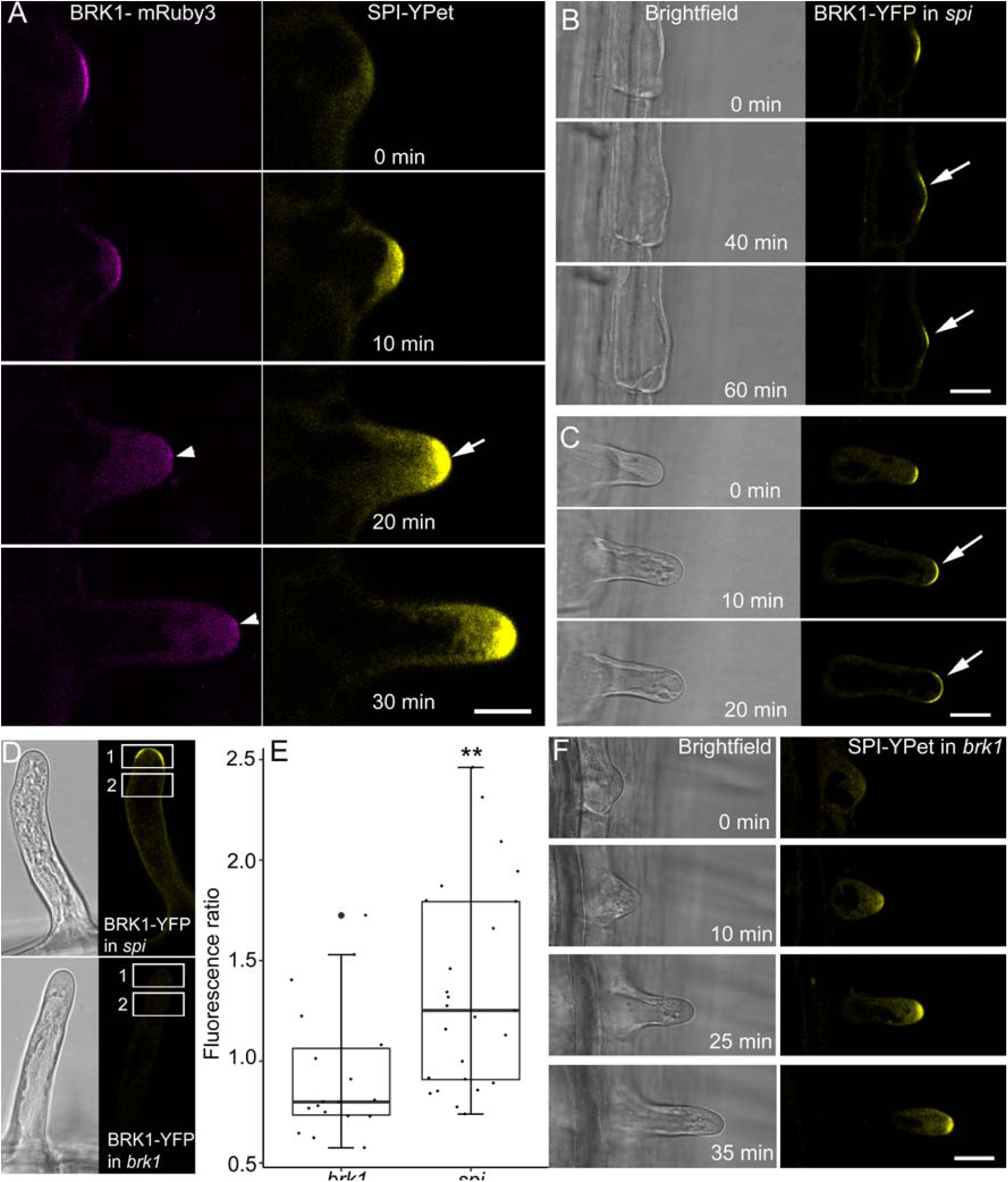
Depletion of BRK1 as Root Hairs Transition to Tip Growth is Delayed in *spi*. **(A)** Dual imaging of BRK1-mRuby3 (arrowheads) and SPI-YPet shows that dissipation of BRK1 coincides with accumulation of SPI (arrow) as root hairs transition to rapid tip growth. Bar = 20 μm **(B)** and **(C)** BRK1-YFP signal persists in *spi* root hair bulges (arrows in B) that fail to transition to tip growth and in slow-growing *spi* root hairs (arrows in **C**). Bars = 20 μm. **(D)** Method for quantification of BRK1-YFP signal persistence in the root tip of *spi*. Root hairs of *brk1* and *spi* expressing BRK1-YFP that were about 40 μm in length were selected. Rectangular region of interests (1 and 2) were drawn to obtain fluorescence values. Ratio values were obtained by dividing mean fluorescence in rectangle region 1 over region 2 used to plot data in panel **E**. **(E)** Box plot of BRK1-YFP root hair tip gradient expressed as fluorescence ratio. Box limits indicate 25^th^ and 75^th^ percentiles, horizontal line is the median and whiskers display minimum and maximum values. Asterisk (**) indicates statistical significance (*p*<0.01) as determined by Student’s T-test. Each dot represents measurement from 3-6 root hairs per group from 8 plants. Only root hairs of similar length were compared. The average root hair length for *brk1* was 37.28 µm (standard deviation 18.05 µm) and for *spi5* was 38.74 µm (standard deviation 18.34 µm). **(F)** Accumulation of SPI-YPet at the root hair tip is not altered in *brk1*. Bar = 20 μm

## DISCUSSION

Our work uncovers new insights underlying actin-mediated root hair development. A major result from our studies is the revelation that SPI is a root hair tip-localized protein. Although SPI fused to fluorescent proteins was reported to localize to endosomes and P-bodies, such studies have been limited to transient expression assays in biolistically-bombarded leaves (Steffens et al., 2015; Steffens et al., 2017). As such, a mechanistic link between reported SPI subcellular localization patterns and cell growth phenotypes (i.e. short root hairs and distorted trichome shapes) described in the original *spi* mutant alleles has never been demonstrated. The large size of SPI and the possibility that it is targeted to discrete cellular domains could have hindered transgenic complementation and subsequent *in planta* localization efforts. Through our work, we have addressed this knowledge gap and now show that a SPI-YPet fusion generated through recombineering has complemented *spi* (Brumos et al., 2020, Zhou et al., 2011). In doing so, we provide compelling evidence that SPI is a root hair tip–enriched BEACH domain-containing protein.

Our studies indicate that SPI mediates root hair tip growth by maintaining the tip-focused fine F-actin meshwork, although the precise mechanism by which this is accomplished is unknown. The root hair tip-focused F-actin meshwork is the functional equivalent of domain-specific actin structures found in other tip-growing cells such as cortical actin fringes in pollen tubes and actin spots in moss protonemata (Bascom et al., 2018a). Although the tip-focused F-actin meshwork was able to form in a population of *spi* root hairs, it often dissipated or was replaced by thick actin cables, a feature of mature wild-type root hairs that are terminating tip growth. Although the observation that the presence of tip-focused SPI-YPet was strongly correlated with maintenance of the tip-focused F-actin meshwork, direct causal links between SPI and F-actin have yet to be established. An alternative explanation is that tip growth induced reduction due to the absence of SPI could lead to downstream, indirect effects on actin organization. The root hair phenotypes and corresponding depletion of tip-focused F-actin in *spi* is reminiscent of studies of BEACH domain-containing proteins in mammalian cells, particularly in neurons, which have often been compared to tip-growing plant cells (Baluška, 2010). For instance, neuronal dendritic spines are cytoplasmic protrusions that modulate excitatory synaptic transmission in the mammalian nervous system. Actin is a major component of dendritic spines and it plays a central role in driving the dynamic shape changes and secretory activities of these cytoplasmic protrusions during synaptic signaling (Cingolani and Goda, 2008). In animals, neurobeachin (Nbea) is a BEACH domain-containing protein that has a similar domain architecture as SPI. Cultured neurons of *Nbea* mice knockouts had fewer dendritic spine protrusions and depleted F-actin at the synapse (Niesmann et al., 2011). Another BEACH domain-containing protein called FAN was shown to be crucial for the formation of filopodia, actin-rich plasma membrane extensions that enable motile cells to probe their environment (Mattila and Lappalainen, 2008). FAN deficient fibroblasts had reduced filopodia formation and were unable to reorganize their actin cytoskeleton in response to upstream activation by tumor necrosis factors (Haubert et al., 2007). Taken together, these results indicate that actin-mediated regulation of cell polarity by BEACH domain-containing proteins is likely to be conserved across animals and plants.

The formation of SPI-YPet agglomerates in response to BFA indicates that SPI is associated with post-Golgi vesicles. This result is consistent with observations made with BEACH domain-containing proteins in mammals. For example, Nbea was shown to localize to vesicular endomembranes adjacent to the *trans*-Golgi network, and like SPI, the distribution of Nbea-positive post-Golgi vesicles was altered by BFA (Wang et al., 2000). Furthermore, the *Caenorhabditis elegans* Nbea homolog, SEL-2, was demonstrated to function in endomembrane traffic in polarized epithelial cells based on the finding that *sel-2* mutants mistarget proteins normally found in the apical cell surface to the basolateral surface (de Souza et al., 2007). Here, the protein secretory marker SEC-RFP, which is typically trafficked to the tips of elongating wild-type root hairs (Sparks et al., 2016), was uniformly distributed in *spi* root hairs. The absence of tip-directed SEC-RFP gradients in *spi* root hairs shows that, as in mammals, plant BEACH domain-containing proteins are required for proper targeting of molecular cargo to points of polarized cell growth.

Another significant result from our studies is the discovery that BRK1 and SCAR2 are molecular determinants of the RHID. The localization of BRK1 and SCAR2 to the RHID and their dissipation as root hair tip growth commenced, contrasts with findings in *Physcomitrella* in which a functional BRK1-YFP distinctly labeled the tips of caulonemal cells (Perroud and Quatrano, 2008). Like BRK1-YFP, an ARPC4-GFP construct labeled the tips of caulonemal cells (Perroud and Quatrano, 2008) whereas no clear tip-focused labeling of ARPC5-GFP was observed in root hairs (Supplemental Figure 5). The observation that BRK1-YFP prominently localized to the RHID, but dissipated during active root hair tip growth was surprising given that the *Arabidopsis BRK1* complemented the defective filamentous growth of *Physcomitrella BRK1* knockouts (Perroud and Quatrano, 2008). *Physcomitrella BRK1 and ARPC4* mutants have clear tip growth defects, while *Arabidopsis* mutants in the *W/SC* and *ARP2/3* complexes have only weak to no tip growth abnormalities (Le et al., 2003, Mathur et al., 2003, Perroud and Quatrano, 2006, Perroud and Quatrano, 2008). This suggests that seed plants might have evolved a specialized function for the W/SC complex at the site of root hair emergence and early bulge formation with only a minor role in driving actin-dependent tip growth. Moreover, a recent discovery regarding the emergence of an ARP2/3 independent W/SC pathway in the regulation of F-actin dynamics for sperm nuclear migration in *Arabidopsis* pollen tubes further indicates that this occurrence may be more common in seedling plants. On the other hand, the knockouts of an *Arabidopsis SPI* orthologue in the liverwort, *Marchantia polymorpha*, led to short rhizoids indicating conserved functions of SPI across land plants in regulating tip growth (Honkanen et al., 2016). In other single cell types such as diffusely-growing trichomes, W/SC and ARP2/3 subunits likely play a more prominent role than SPI based on their more severe trichome phenotypes when expression of their encoding genes are suppressed.

The weakening of BRK1-mRuby3 fluorescence coinciding with SPI-YPet accumulation, and the persistence of SPI-YPet signal in *spi* root hair tips, provides indirect evidence that SPI might play a role in mediating BRK1 stability or localized clustering at the plasma membrane of the RHID. Although, it is unknown why BRK1-YFP signal persists in *spi*, it is tempting to speculate that SPI might modulate BRK1 via protein degradation pathways. This possibility is supported by studies in mammals pointing to a role for BEACH domain-containing proteins in protein degradation via the ubiquitination pathway. In mouse models for example, the BEACH domain-containing protein, WDR81, was shown to be essential for removal of autophagy-dependent ubiquitinated proteins (Liu et al., 2017). In this regard, it is worth noting that the W/SC-ARP2/3 pathway was demonstrated to function in stress-induced autophagy (Wang et al., 2016) and proteasome inhibitors stabilized SCAR during dark-induced primary root growth inhibition (Dyachok et al., 2011). It is possible that SPI-mediated proteolytic pathways and BRK1 might coordinate their activities to specify the levels of W/SC at the RHID that enables the transition to actin-dependent rapid tip growth. However, such scenario is complicated by the observation that the formation of tip-directed SPI-YPet does not appear to require BRK1. This suggest that SPI’s appearance at the root hair tip is likely regulated by other factors besides W/SC complex. Alternatively, the failure of BRK1 to disappear from the RHID in *spi* mutants may be caused by general failure to transition to normal tip growth. Future studies will require subjecting root hairs to conditions that prematurely terminate tip growth to determine if BRK1 signals persist.

In summary, our work provides new data that contribute to our understanding of actin-mediated root hair development. A crucial result from our work is the discovery that SPI and the W/SC subunits, BRK1 and SCAR2, exhibit polarized localization patterns in root hairs that point to potential functional relationships among these proteins during root hair development. For the future, it will be important to evaluate genetic interactions between SPI and W/SC, and ask whether SPI physically interacts with actin or subunits of W/SC to better explain the functional links between SPI and W/SC.

## METHODS

### Forward-Genetic Screening and Map-Based Cloning

The *hlb* mutants from which the *spi-5* mutant allele was identified were isolated from a population of Transfer-DNA (T-DNA) seed (Arabidopsis Biological Research Center stock CS31100). Plants from which the seeds were derived from were transformed with the activation tagging pSKI015 plasmid (Weigel et al. 2000; Sparks et al., 2016; Sun et al., 2019). Briefly, mutagenized seeds were surface sterilized in 95 % (v/v) ethanol and 20% bleach (v/v), followed by three washes in sterile deionized water. A solution of 0.5 x Murashige and Skoog (MS) basal salt medium with vitamins (PhytoTech Labs, USA) and 1 % (w/v) sucrose was prepared, and the pH of the solution was adjusted to 5.7. After adding 0.5% agar (w/v) (Sigma-Aldrich), the solution was autoclaved and allowed to cool to room temperature. Upon reaching 55°C, a stock solution of 10 mM LatB (CalBiochem-EMD Chemicals) in 100% Dimethyl Sulfoxide (DMSO) was added to make a final LatB concentration of 100 nM. Sterilized seeds were suspended in the MS-agar-LatB medium and gently swirled to evenly distribute the seed. The seed-MS-agar-LatB mixture was poured to a thickness of 2 mm on the base of 10 cm x 10 cm Petri dishes. After keeping plates at 4°C for 48 h, they were positioned vertically in a Conviron set to 24°C with a 14 hours light (120 µmol m^-2^s^-1^)/ 10 hours dark cycle. Six days after transfer to the Conviron growth chamber, seedlings that exhibited severe growth inhibition were transplanted to LatB-free media and grown to maturity.

For LatB hypersensitivity assays, seed from selected plants and the *hlb2* mutant were planted on the surface of a 3 mm layer of polymerized 0.5 x MS - 1% agar growth media in gridded square 10 cm × 10 cm Petri plates and grown in the same Conviron used for screening. Four-day-old seedlings with primary roots that were about 1 cm long were selected. Selected seedlings were transplanted to a new set of square Petri dishes containing MS medium supplemented with 50 nM LatB or MS supplemented with the appropriate volume of LatB solvent (i.e. DMSO). During transplant, the tip of the root was positioned at the grid line of the Petri dish and maintained in a vertical orientation. Four days after transplanting, images of the roots were captured with a Nikon Insight digital camera mounted on a copy stand. Primary root length was expressed as the distance between the position of the root tip 4 days after transplant and the grid line where the root tip was positioned during transplant (Supplemental Figure 1).

To identify the *HLB2* gene, homozygous *hlb2* (Col-0 ecotype) was out-crossed to the Landsberg *erecta* ecotype to generate seeds for map-based cloning because attempts to identify the responsible mutation for *hlb2* phenotype using TAIL-PCR had been unsuccessful. Segregating F2 seedlings were surface sterilized as described above and grown for three days. These seedlings were then transferred to MS media containing 50 nM LatB and root lengths were marked on the plates to track root growth. Transferred seedlings were grown vertically for an additional four days in the growth chamber and seedlings that showed root hypersensitivity to 50 nM LatB were selected for mapping. Briefly, DNA was extracted (Edwards et al., 1991) from about 2600 LatB sensitive seedlings and cloned with simple sequence length polymorphism (SSLP) and cleavage of amplified polymorphic site (CAPS) markers to chromosome one between *AT1G02740* and *AT1G03410* loci (Lukowitz et al., 2000), which spanned 248 kb with 77 annotated genes. Primers were then designed to several candidate genes based on the sequenced 248 kb region. Nucleotide sequencing revealed that *hlb2* mutation had a 10 base pair deletion (Chr1 position 720,152-720,161) in exon 14 of the *AT1G03060* gene. In addition, we obtained a T-DNA insertional mutant (SALK_065311) from the ABRC with predicted insertion in *SPI*. After genotyping, SALK_065311 was subjected to similar LatB hypersensitivity and growth assays as *hlb2* as aforementioned. Allelism was determined by examining the F1 progeny from a cross between *hlb2* × SALK_065311. Following the nomenclature of Steffens et al. (2015), SALK_065311 and *hlb2* were named *spi-3* and *spi-5*, respectively.

### Generation of Fluorescent Protein-Tagged Constructs and Plant Lines

The SPI protein was tagged at the C-terminus with the fluorescent protein 3x-YPet by the recombineering method (Zhou et al., 2011). *Agrobacterium tumefaciens* UIA143 pMP90 harboring a SPI-YPet recombineering construct was used to transform the *spi-5* mutant by the floral dip method (Clough and Bent, 1998). Transgenic lines were isolated based on resistance to kanamycin and restoration of wild-type root length. Presence of the SPI-YPet transgene was amplified with primers (5’-ATTCCACAAGCAACCAGTCAC-3’ and 5’-AACAGAGTTGAGAGTGGCTCG-3’) and sequenced to confirm the correct configuration. Final validation of *SPI-YPet* expression was accomplished by screening selected lines under the confocal microscope for YPet fluorescence and complementation of the short root hair phenotype.

*SCAR2-mCherry* lines were also generated by recombineering in which an mCherry tag was fused to an internal region of SCAR2 (Sharan et al., 2009). The mCherry tag was inserted after amino acid 585 in SCAR2 to generate SCAR2-i2mCherry. Recombineering primers were designed as follows: the forward primer contained 50 SCAR2 nucleotides upstream from the insertion site of SCAR2 and 18 nucleotides of the 5’ end of the mCherry cassette, which had a 5x Glycine, 1 Alanine linker. The reverse primer contained 50 nucleotides after the insert site of SCAR2 and 24 nucleotides of the 3’ end of the mCherry cassette. The JAtY75L14 clone was transformed into the recombineering competent *E.coli* strain SW105 and recombined with the SCAR2-i2mCherry recombineering cassettes. A flippase recombination reaction removed the ampicillin resistance marker. The sequence between the two test primers was verified by DNA sequencing. The clones were transformed into *Agrobacterium* and then into *scar1234* plants (Dyachok et al., 2008). Because both *scar1234* plants and the recombineering clones were Basta resistant, plants on MS plates were screened by eye for rescued trichome phenotypes. Rescued plants were genotyped with the recombineering test primers and screened for fluorescence with a confocal microscope.

The *BRK1promoter: BRK1-YFP* introduced into the *brk1-1* mutant and *ARPC5-GFP* are described in Dyachok et al. (2008) and Yanagisawa et al. (2015), respectively. For the *BRK1-mRuby3* construct, the fluorescent protein 3x-mRUBY3 was tagged with a C-terminal linker (10 Alanine, Glycine) using Thermo Fisher Scientific GeneArt to include BamH1 and XbaI sites at its 5’ and 3’ ends, respectively. Codon optimization for *Arabidopsis* was performed on the 3x-mRUBY3 and internal linkers prior to synthesis. The 3X-mRuby3 fragment was inserted as a BamH1/XbaI into the plasmid *pBRK:YFPpEZRK* (Dyachok et al., 2008).The *Agrobacterium* floral dip method (Clough and Bent, 1998) was used to transform *brk1-2* plants, and transformation was confirmed when this construct fully restored defective *brk1-2* trichomes.

Plants expressing *BRK1-YFP* were crossed with *spi-5* to generate *BRK1-YFP* in *spi-5* lines. In parallel, plants expressing *SPI-YPet* were crossed with *brk1* to obtain SPI-YPet in *brk1* lines. *Spi-5* was directly transformed with the *UBQ10:mGFP-Lifeact* construct by the floral dip method (Clough and Bent, 1998). *Spi-5* expressing SEC-RFP was generated by crossing *spi-5* with *SEC7-RFP*-expressing wild-type plants and progeny in subsequent generations that exhibited fluorescence and the *spi* root hair phenotypes were selected for analysis.

### Generation of Dual Fluorescent Protein-Labeled Plant Lines

To generate plant lines co-expressing *SPI-YPet* and *BRK1-mRuby3*, *SPI-YPet* in *spi-5* was crossed with *BRK1-mRuby3* in *brk1-2*. Progeny from subsequent generations that exhibited yellow and red fluorescence and rescued root hair and trichome phenotypes were selected for analysis. For lines expressing *SPI-YPet* and *mRuby3-Lifeact* (Bascom et al., 2018b), *SPI-YPet* in *spi-5* was directly transformed with a *UBQ10:Lifeact-mRuby3* construct. Seedlings that showed both YPet and mRuby3 fluorescence were selected under the confocal microscope.

### Evaluation of Root Hair Growth Rate

Seeds were surface-sterilized in ethanol and bleach as described above. To evaluate root hair growth rate, seeds were planted on 48 × 64 mm coverslips coated with 0.5 x MS, 1% sucrose and 0.4 % (w/v) Gelzan ^TM^ CM (Sigma Aldrich, USA) according to Dyachok et al. (2016). Coverslips were placed in 9 cm round Petri dishes and stratified in 4°C for 48 h. After stratification, the coverslip system with planted seed were kept in a 24°C growth chamber with 14 hours light (120 µmol m^-2^s^-1^)/ 10 hours dark cycle for 5 to 6 days.

To quantify root hair growth rate, time lapse sequences of elongating root hairs at intervals of 10 mins for 60 mins from a region of the primary root located between 2 to 3 mm from the root tip were captured with a Nikon Eclipse TE300 inverted microscope using a 10x objective. Root hair lengths at various time points were extracted from digital images using ImageJ (v1.51) software (https://imagej.nih.gov/ij/). The displacement of the root hair tip after each 10 min interval was obtained and divided by 10 to obtain growth rate as μm/min.

### Chemical treatments on root hairs

For BFA treatment, a stock solution of 10 mM was made by dissolving BFA powder (Sigma-Aldrich) in DMSO. Subsequently, the working solution of 50 µM was diluted in 0.5 x MS, 1% sucrose solution and loaded into a 1 mL syringe. The BFA solution was injected directly next to the roots of 4-5 days old *Arabidopsis* seedlings grown on coverslips and was incubated in room temperature for 10 minutes prior to microscopy.

FM 1-43 dye (ThermoFisher Scientific) was dissolved in DMSO to make a 10 mM stock solution. The dye was diluted to 4 µM in 0.5 x MS, 1% sucrose solution. The dye was loaded into a 1 mL syringe and the solution was injected next to the roots as described above.

### Microscopy and Image Analysis

Live cell imaging of root hairs using confocal microscopy was performed on 4 or 5-day-old seedlings grown on the 48 mm × 64 mm coverslip system described above. Coverslips with the seedlings were placed horizontally on the stage of an inverted Leica SP8-X point scanning confocal microscope (Leica Microsystems, Buffalo Grove, Illinois) or an UltraView ERS spinning-disc confocal microscope (Perkin Elmer Life and Analytical Sciences, Waltham, Massachusetts) equipped with 40 × water (numerical aperture=1.10) or 100 × oil (numerical aperture=1.40) immersion objectives. SPI-YPet and YFP-BRK1 were imaged by illuminating roots growing along the coverslip surface with the 514 nm line of the SP8-XArgon laser and emission detected at 527 nm. Images of root hairs expressing SCAR2-mCherry, BRK1-mRuby3, Lifeact-mRuby and SEC7-RFP were acquired through illumination with the tunable SP8-X white light laser (560-580 nm) and detecting emission at 610 nm. Excitation and emission parameters for GFP (GFP-ARPC5 and mGFP-Lifeact) were 488 nm and 510 nm, respectively. Time-lapse movies or single time point images were collected using Volocity acquisition version 6.3.5 (Improvision) or SPX-8 LAS software, for the UltraView and Leica SP8-X, respectively.

Quantification of fluorescence from root hair images was conducted on 8-bit confocal images acquired at a single fixed focal plane that spanned the median of the cell (SPI-YPet, BRK1-YFP, SEC-RFP, and BFA and FM 1-43 treatments) or from maximum projected images (mGFP-Lifeact). For SPI-YPet and SEC-RFP, an oval region of interest (ROI) at the root hair tip was drawn and mean fluorescence within this area was acquired using Image J. Fluorescence was expressed as the ratio of mean fluorescence within the root tip ROI to background fluorescence. For SPI-YPet, the background used was the region adjacent, but outside the root hair tip (Figure 1C) while for SEC-RFP, the background used was an area on the sub-apical region of the root hair tip (Figure 2F). For BRK1-YFP, a ROI was drawn along the apical-most root hair tip that was about 20 pixels-wide using the selection brush tool of Image J. The ratio of fluorescence within this area to background fluorescence was obtained (Figure 2C). For BFA, a variable ROI was selected based on the size of BFA-induced agglomerates using the selection brush tool of Image J. The ratio of fluorescence was calculated based on similarly sized ROI in the cytoplasmic region. For FM 1-43 experiment, the same quantification technique as SEC-RFP was used.

Root hair growth rate data for scatter plots in Figures 1 and 5 were derived from the same root hair images in which fluorescence images were acquired on Leica SP8-X. For the former, an image of a root hair was taken at time 0 and every 5 minutes thereafter. Scatter plots to determine the relationship between growth rate and tip-focused fluorescence was done in R (R Core Team, 2019) using ggplot function in ggplot2 package (Wickham, 2016). Linear regression analysis between two variables were performed using lme4 package on R software (Bates et al., 2015, R Core Team, 2019).

Root hair growth rate for Figures 3Q was obtained directly on Nikon TE300 from root hairs growing in 0.5 X MS, 1% sucrose and 0.4% Gelzan in 5 cm diameter round petri dishes. Time lapse images were taken every 10 minutes over a period of 60 minutes. The growth rate refers to the displacement of the root hair tip in μm divided by time elapsed (min).

To quantify tip-focused F-actin, 25 optical sections were taken at 0.5 μm intervals using the UltraView spinning-disc confocal microscope. Raw Ultraview Z-stacks were exported to Imaris image analysis software version 9.2.0 (Bitplane). Transverse sections of the root hair tip were obtained from Z-stacks using the surpass view interface of the Imaris software and exported as 8-bit TIFF files. From these images, an ROI spanning the circular area of the root hair tip was drawn using image J and mean fluorescence was extracted (Figure 3O). The ratio of the tip fluorescence to background was obtained from 18-21 root hairs.

Violin plots for planar polarity in Figure 5F were designed with ggplot function in ggplot2 package (Wickham, 2016). Pairwise Kolmogorov-Smirnov test was selected to compare the shape of two empirical cumulative distributions for planar polarity of two genotypes using basic package in R (R Core Team, 2019).

## Accession Numbers

Sequence data from this article can be found in the Arabidopsis Genome Initiative or GenBank/ EMBL databases under the following accession numbers: *SPI* (AT1G03060), *BRK1* (AT2G22640), *SCAR1* (AT2G34150), *SCAR2* (AT2G38440), *SCAR3* (AT1G29170), and *SCAR4* (AT5G01730)

## Supplemental Data

**Supplemental Figure 1.** Primary Root Growth of *hlb2* Seedlings is Hypersensitive to LatB.

**Supplemental Figure 2.** H*L*B2 Encodes the Beach Domain Containing Protein, SPIRRIG.

**Supplemental Figure 3.** S*P*I*-YPet* Construct Complements *spi*

**Supplemental Figure 4.** FM1-43 Assays in Growing wild-type and *spi* Root Hairs

**Supplemental Figure 5.** ARP2/3 Pathway Does Not Mark the Root Hair Initiation Domain

**Supplemental Movie 1**. Time-Lapse Confocal Microscopy of SPI-YPet in Multiple Elongating Root Hairs

**Supplemental Movie 2**. Time-Lapse Confocal Microscopy of SPI-YPet in a Single Rapidly Elongating Root Hair.

**Supplemental Movie 3**. Time-Lapse Confocal Microscopy of SPI-YPet in a Root Hair Bulge Transitioning to Tip Growth.

**Supplemental Movie 4**. Time-Lapse Confocal Microscopy of SPI-YPet in a Root Hair during Termination of Tip Growth.

**Supplemental Movie 5**. Spinning-Disc Confocal Microscopy of Lifeact-mGFP in Elongating Root Hairs of Wild type and *spi*.

**Supplemental Movie 6**. Spinning-Disc Confocal Microscopy of a Rapidly Elongating Root Hair Co-Expressing SPI-YPet and mRuby3-Lifeact.

**Supplemental Movie 7**. Time-Lapse Confocal Microscopy of BRK1-YFP Root Hair Bulge Transitioning to Tip Growth.

**Supplemental Movie 8**. Time-Lapse Confocal Microscopy of a Root Hair Bulge Transitioning to Tip Growth and Co-Expressing SPI-YPet and mRuby3-Lifeact.

**Supplemental Movie 9**. Time-Lapse Confocal Microscopy of a *spi* Root Hair Expressing BRK1-YFP.

## ACKNOWLEDGMENTS

We thank Drs. Jose Alonso, Linda Robles and Anna Stepanova (North Carolina State University) for assistance with recombineering and Dr. Magdalena Bezanilla (Dartmouth College) for the Lifeact-mRuby construct. This work was supported by the National Aeronautics and Space Administration (NASA grants 80NSSC19K0129 and 80NSSC18K1462) and the Noble Research Institute to E.B.B.) and by the National Science Foundation (NSF MCB Grant No.1715544) to D.B.S. We also thank Dr. Larry M. York (Noble Research Institute) and Dr. Wayne Versaw (Texas A.M. University) for critical comments on the manuscript.

## AUTHOR CONTRIBUTIONS

S.C., T.K, D.B.S and E.B.B. conceptualized and designed the research, and analyzed and interpreted the data. S.C., J.A.S. and E.B.B. conducted root hair, primary root growth, microscopy, image and statistical analysis of data. T.K., J.A.S., B.R.K., E.L.M. and D.B.S. generated various research reagents. All authors contributed to writing the manuscript.

